# NanoSquiggleVar: A method for direct analysis of targeted variants based on nanopore sequencing signals

**DOI:** 10.1101/2023.03.15.532860

**Authors:** Jidong Lang

**Affiliations:** Department of Bioinformatics, Qitan Technology (Beijing) Co., Ltd, Beijing, China

**Keywords:** nanopore sequencing technology, squiggle segment, variation calling, dynamic time warping (DTW), long-read sequencing

## Abstract

**Background:** Nanopore sequencing is a fourth-generation sequencing technology that has developed rapidly in recent years. It has long sequencing read lengths and does not require the polymerase chain reaction to be performed. These characteristics give it unique advantages over the next-generation sequencing technology under certain usage scenarios. The number of bioinformatics analysis algorithms and/or tools developed with nanopore sequencing has increased sharply during the past years, undoubtedly providing great help and support for the application of nanopore sequencing in scientific research and practical scenarios.

**Results:** We developed NanoSquiggleVar, a method for direct analysis of targeted variants based on nanopore sequencing signals. It first establishes a set of wild-type and mutant-type target signals within the same experimental and sequencing system, named wild squiggle set and variant squiggle set, respectively. In each sequencing iteration, the signal is sliced into fragments by a moving window of 1-unit step size. Then, dynamic time warping is used to compare the signal squiggles to the detected variants. Point mutations, insertions and deletions (indels), and homopolymer sequences were simulated and generated by Scrappie and then analyzed and evaluated with NanoSquiggleVar. We found that all of these variants were efficiently detected and discriminated, and the results were consistent with the expectations.

**Conclusions:** NanoSquiggleVar can directly identify targeted variants from the nanopore sequencing electrical signal without the requirement of base calling, sequence alignment, or variant detection with downstream analysis. We hope that this method can complement targeted variant detection using nanopore sequencing and potentially serve as a reference for real-time sequencing and analysis.

## INTRODUCTION

Nanopore sequencing, also known as fourth-generation sequencing or single-molecule real-time DNA sequencing, identifies single DNA molecules without requiring the polymerase chain reaction (PCR). No PCR amplification or chemical labeling is involved in the real-time sequencing of DNA or RNA molecules through nanopore sequencing, thus avoiding the introduction of false mutations in the process and ensuring high accuracy. The current next-generation sequencing (NGS) technology is capable of generating reads of hundreds of bases, whereas nanopore sequencing produces reads of several kilobases and even ultra-long reads of several megabases (Magi et al., 2018; Wang et al., 2021). Nanopore sequencing is performed by moving single-stranded DNA/RNA through a membrane via nanopores and applying voltage across the membrane. Nucleotides present in the pores affect the electrical resistance at those loci, and the current measured indicates the DNA/RNA base sequence passing through the pores over time. The current signal (plotted as a squiggle) is the raw data collected by the nanopore sequencing instrument (Rang et al., 2018; Wick et al., 2019).

Base calling is the conversion of the original signal generated by nanopore sequencing into base sequences. This is typically not a simple task because the electrical signal coming from individual molecules can generate noise and random data. Studies have compared the base calling methods currently in use on the ONT sequencing platform, such as Guppy (https://community.nanoporetech.com), Scrappie (https://github.com/nanoporetech/scrappie), Bonito (https://github.com/nanoporetech/bonito), and Chiron (Teng et al., 2018), and found that different methods and parameters have different performance on data. The robustness of these base calling methods could also fall short on species that are different from the training data of the existing models (Wan et al., 2022; Wick et al., 2019). As a result, it is often necessary to perform species-specific model training to maximize the accuracy of base calling. Larger neural networks also give more accurate results, including a significant improvement on reading accuracy, but they require higher computational efficiency.

An increasing number of bioinformatic analysis methods and tools are being developed for nanopore sequencing data for different application scenarios, such as the alignment tool Minimap 2 (Li, 2018), mutation detection tools PEPPER-Margen-DeepVariant (Shafin et al., 2021) and Nano2NGS-Muta (Lang et al., 2022), genome assembly tools Canu (Koren et al., 2017) and FlyE (Kolmogorov et al., 2019), and short tandem repeats detection tools Tandem-genotypes (Mitsuhashi et al., 2019) and NanoSTR (Lang et al., 2023). However, all these tools depend on the accuracy of base calling in their data conversion. More importantly, the different approaches and principles of the tools as well as the principles and data characteristics of nanopore sequencing (Magi et al., 2017; Wang et al., 2021) inevitably result in their varied performance in variant detection. This poses a great challenge for users in their selection of data analysis tools.

To tackle these problems, we developed NanoSquiggleVar, a method for direct analysis of targeted variant detection from nanopore sequencing signals. It eliminates base calling, sequence alignment, and variant calling in the downstream data analysis, thus enabling direct identification of the targeted variations from nanopore sequencing signals. NanoSquiggleVar avoids the problems associated with base calling of nanopore sequencing for different species and saves the cost of the large-scale training data required to build the base calling models. It also reduces the impact of different data analysis methods on variant detection in the downstream data analysis.

## METHODS AND MATERIALS

### Principle of NanoSquiggleVar

As shown in Figure 1, NanoSquiggleVar analysis involves five steps: i) establish the wild squiggle set (WSS) and variant squiggle set (VSS) for variant targeting; ii) measure the nanopore sequencing signal; iii) based on the length of each squiggle set (fragment size), slice the current signal into squiggle segments using a moving window of 1-unit step size; iv) use dynamic time warping (DTW) to align the sliced squiggle with both the WSS and VSS, and calculate the respective distance values, namely WSS-Distance and VSS-Distance; v) analyze the WSS-Distance and VSS-Distance, and retain data that satisfy the following three criteria: WSS-Distance <= threshold, VSS-Distance <= threshold (for example, the reference threshold for Scappie-simulated data was set at 18), and |WSS-Distance − VSS-Distance| > 1. Define Score is the average value of WSS-Distance divided by VSS-Distance, with Score > 1 indicating variant type and Score <= 1 indicating wild type.

**Figure 1.**
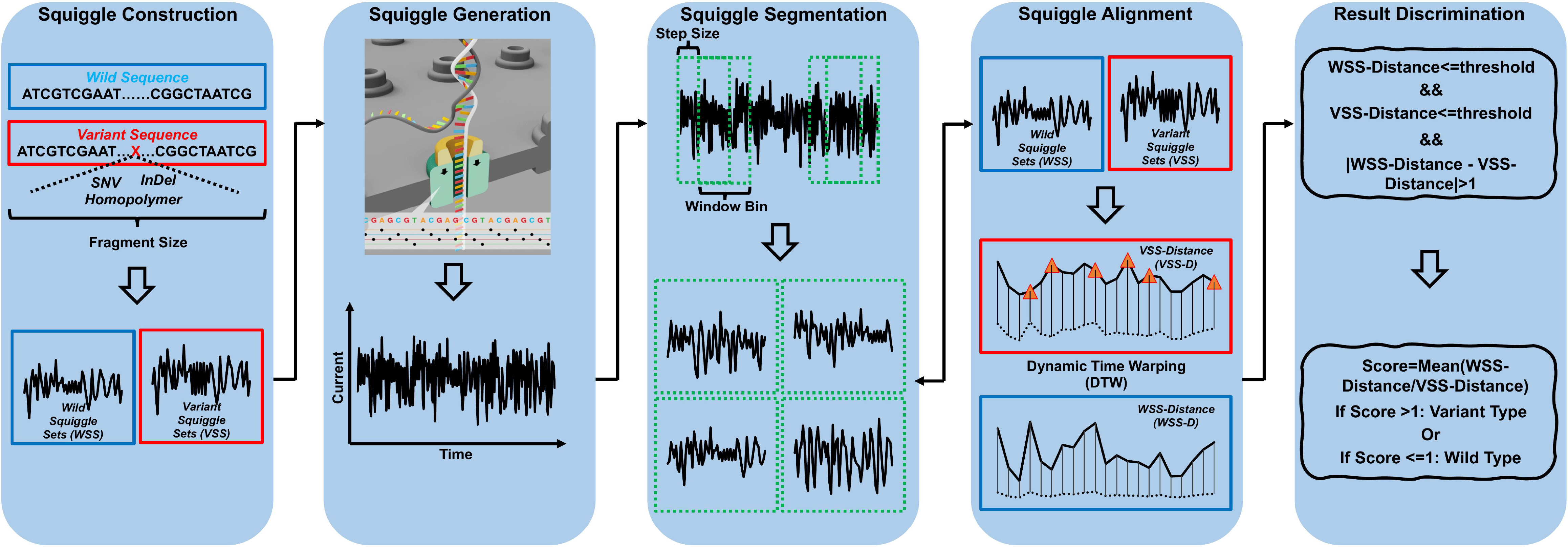
Principle of NanoSquiggleVar analysis.

### Simulation of nanopore sequencing signals with Scrappie

The genome sequences of six different viruses were simulated. With the SARS-CoV-2 reference genome MN908947.3 as the template, the four unique mutation sites of the alpha, beta, delta, gamma, lambda, and omicron variants were used for point mutation substitutions at the corresponding loci (Lang, 2022) (Supplementary Sheet Table S1), which provided six reference sequences containing different mutation sites in addition to the reference genome without mutations. Ten 1–10 bp bases were randomly inserted into the reference genome sequence, and 10 1–10 bp bases were randomly deleted. Three A-bases with the lengths of 10 bp, 15 bp, and 20 bp were also randomly inserted into the reference genome sequence. The above steps were performed to simulate indels and homopolymer sequences (Supplementary Sheet Table S2) in the genome. All of these mutation sites were taken as the centers to extract 30 bp from the upstream side and downstream side as the fragment sequences. Scrappie *squiggle* was used to transform the fragment sequences and reference sequences into the nanopore sequencing signals, that is, the squiggle set.

### Simulation of nanopore sequencing data

Deepsimulator (version v1.5) (Li et al., 2018; Li et al., 2020) was used to simulate the nanopore sequencing data of seven genome sequences. Minimap2 (version 2.21-r1071) and Sambamba (version 0.8.0) (Tarasov et al., 2015) were used to align the simulated data to six reference sequences with mutations and sort the SAM files. Fifty mapped reads with mutations were randomly selected and mixed with 950 randomly selected reads from data simulated by Seqkit (version v0.16.1) (Shen et al., 2016) using reference sequences without mutations to generate the final simulated nanopore sequencing data. Longshot (version 0.4.1) (Edge and Bansal, 2019) was used for mutation detection.

### Simulation of NGS sequencing data

Wgsim (version 1.10) (https://github.com/lh3/wgsim) was used to simulate the NGS sequencing data of seven genome sequences. Bwa (version 0.7.17-r1188) (Li and Durbin, 2009) and Samtools (version 1.12) (Li et al., 2009) were used to align the simulated data to six reference sequences containing mutations and sort the SAM files. Fifty mapped reads with mutations were randomly selected and mixed with 950 randomly selected reads from the data simulated by Seqkit (version v0.16.1) from reference sequences without mutations to produce the final simulated NGS sequencing data. Freebayes (version v1.0.2) (Garrison and Marth, 2012) was used for mutation detection.

## RESULTS

### NanoSquiggleVar accurately and effectively detects point mutations

We performed targeted point mutation detection and analysis on Scrappie-simulated signals and found that NanoSquiggleVar effectively detected the expected mutations with a 100% consistency (Supplementary Sheet Table S3). Taking C12534T of Sample-1 as an example, its WSS and VSS squiggle signals were compared with that of the genomes of the simulated data Sample-1 to Sample-6 and the references using DTW. The results showed that for all simulated data, a significant trough existed between position 12482 and position 12547 (Figure 2A), indicating the presence of the C12534T mutant signal and wild-type signal at this location. For C12534T of Sample-1, the DTW distance of the wild-type squiggle was greater than that of the mutant-type squiggle (Figure 2B), corresponding to Score > 1 and therefore indicating the presence of the C12534T mutant signal in Sample-1. This was consistent with our expectation. Among all data-simulated squiggles with the C12534T mutation, the shortest distance was observed between Sample-1 and the genomes (Figure 2C), suggesting the occurrence of the C12534T mutation in Sample-1. Similarly, for the squiggle signal with no C12534T mutation, the distance between Sample-1 and the genomes was the largest (Figure 2D), again indicating the C12534T mutation in Sample-1.

**Figure 2.**
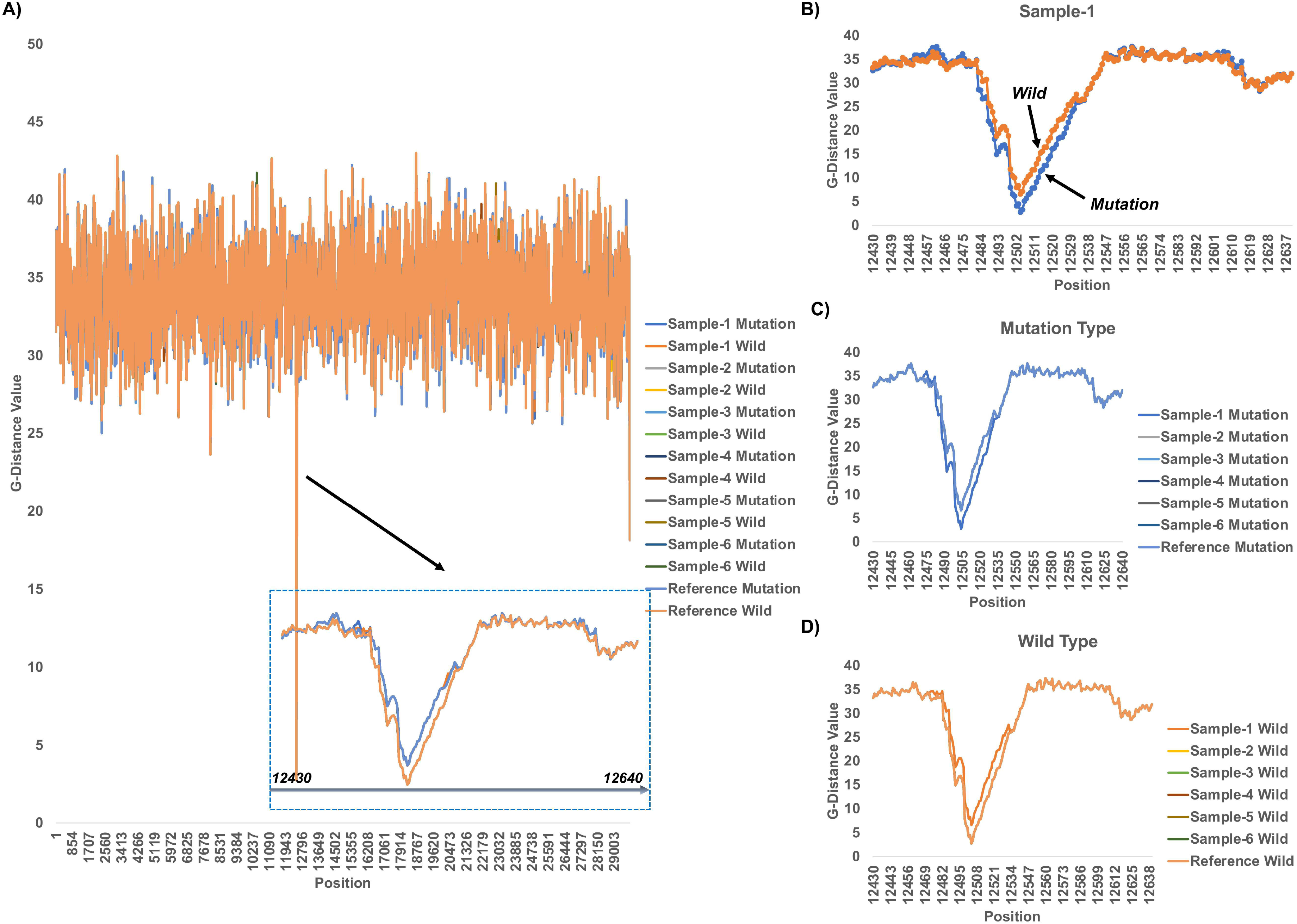
NanoSquiggleVar analysis of C12534T in Sample-1. A) Distance distribution of mutant and wild-type C12534T squiggle on the seven genomes. The plot in the blue dotted frame shows the distribution of the DTW distance at position 12482-12547. B) Distance distribution of wild-type and mutant C12534T squiggle of Sample-1 on genomes with mutations. C) Distance distribution of mutant C12534 squiggle on the seven genomes. D) Distance distribution of wild-type C12534 squiggle on the seven genomes.

### NanoSquiggleVar accurately and effectively detects indels

Targeted indel detection and analysis were performed on the Scrappie-simulated signals. NanoSquiggleVar effectively detected them with 100% consistency. For example, as shown in Figure 3A, we compared and analyzed the WSS and VSS of the targeted insertion site with the current simulated for the reference genome and found that the corresponding signal could be found near the targeted location and it was predicted to be a wild type. Similarly, as shown in Figure 3C, when we compared and analyzed the WSS and VSS of the targeted insertion site with the current simulated for the mutant genome, we again found that the corresponding signal could be found near the targeted location and it was predicted to be a mutant type. The targeted deletion sites of the reference genome and variant genome were also correctly identified (Figure 3B, Figure 3D).

**Figure 3.**
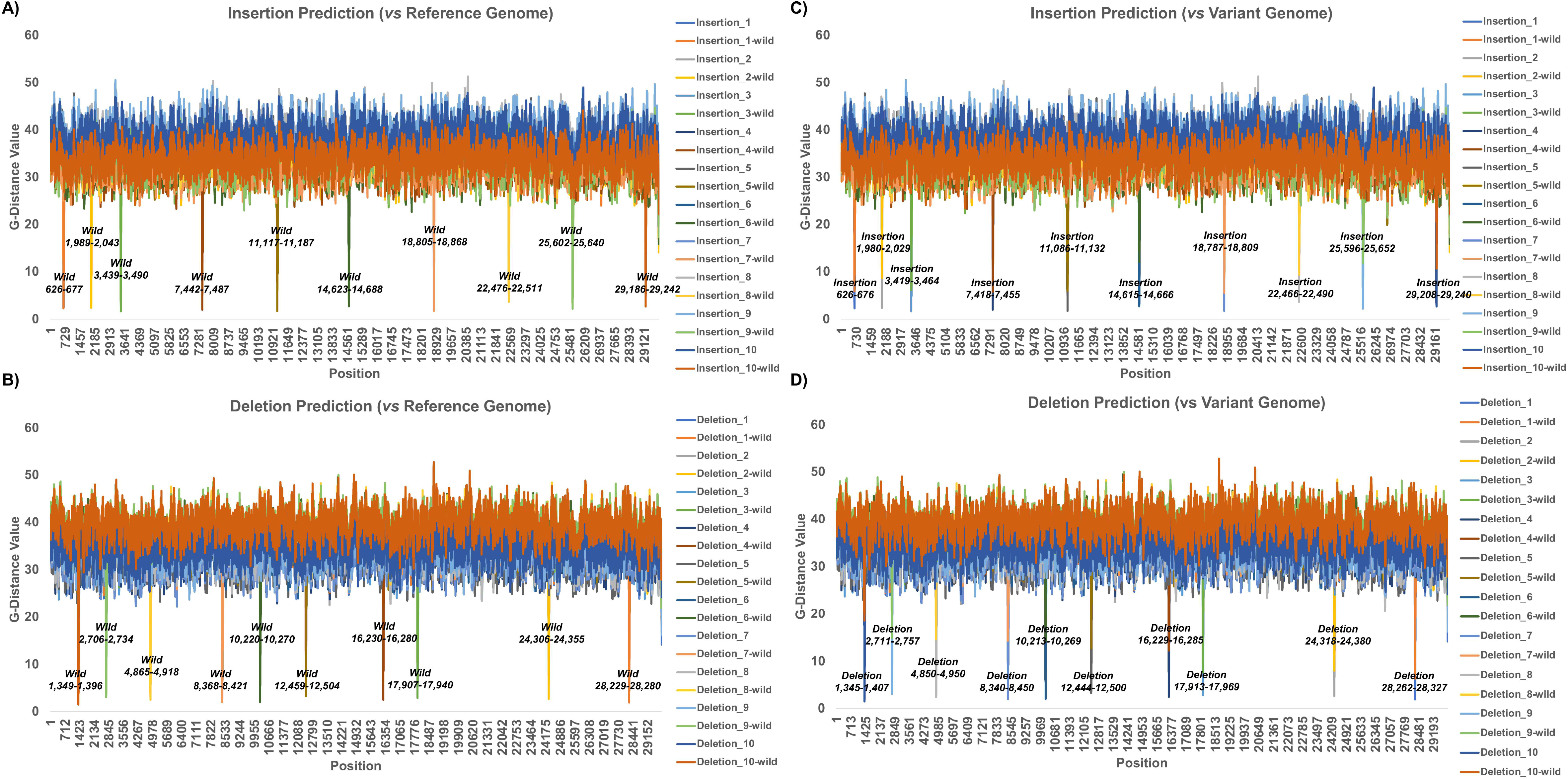
A) Distance distribution of the squiggle alignment between the WSS and VSS of the targeted insertions and the reference genome. B) Distance distribution of the squiggle alignment between the WSS and VSS of the targeted deletions and the reference genome. C) Distance distribution of the squiggle alignment between the WSS and VSS of the targeted insertions and the variant genome. D) Distance distribution of the squiggle alignment between the WSS and VSS of the targeted deletions and variant genome.

### NanoSquiggleVar accurately and effectively detects homopolymer sequences and types

Targeted homopolymer sequence detection and analysis were performed on the Scrapie-simulated signals. NanoSquiggleVar also effectively detected all of the expected homopolymer sequence targets, but the threshold for Score needed to be adjusted from 1 to 0.8. As shown in Figure 4A, we compared the WSS and VSS signals of three homopolymers with the signals of the reference genome and found that the homopolymers could be detected. Notably, different homopolymers exhibited a “step-like” phenomenon, which also showed that our method may be able to effectively distinguish homopolymers from wild types. Similarly, as shown in Figure 4B, we compared the WSS and VSS signals of three homopolymers with the signals of the variant genome and found that the homopolymers could also be detected. Although the “step-like” phenomenon was also present and the correct homopolymer type could be identified, we found that the distance value distribution for multiple A-bases may be mixed with that for the wild type, which interfered with the differentiation of multiple A-bases from the wild type. Therefore, it may be necessary to adjust the Score threshold of NanoSquiggleVar to obtain better results.

**Figure 4.**
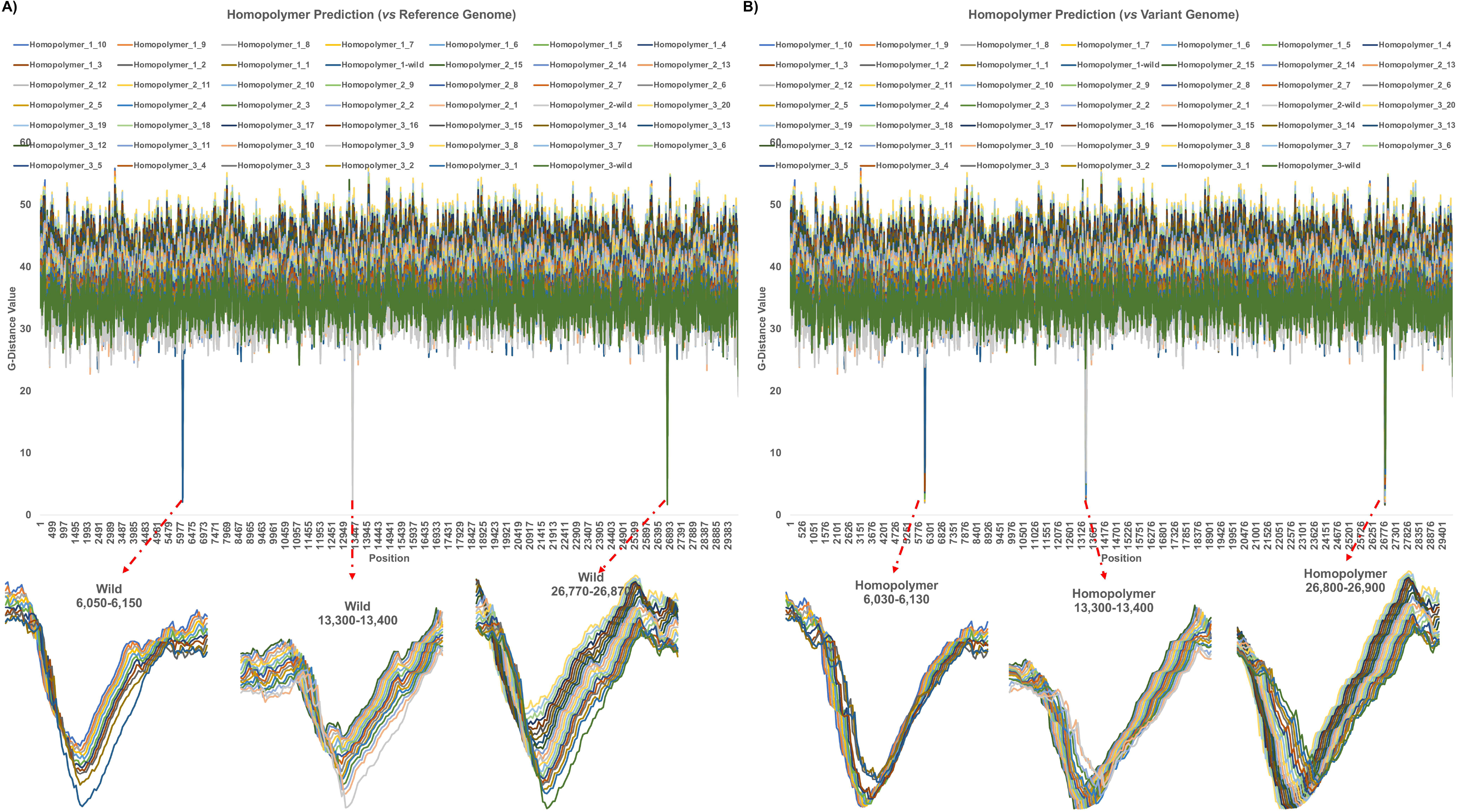
A) Distance distribution of the squiggle alignment of the WSS and VSS of three targeted homopolymer sequences with the reference genome. B) Distance distribution of the squiggle alignment of the WSS and VSS of three targeted homopolymer sequences with the variant genome.

## DISCUSSION

In the past decade, the long-read length and real-time sequencing capability of nanopore sequencing technology made it one of the most effective methods for genome analysis and whole-genome mutation profiling. Examples include the analysis of structural variations, mutations, copy number variations, RNA alternative splicing, and tumor origin tracing (Chen et al., 2020; Goenka et al., 2022; Katsman et al., 2022; Martignano et al., 2021). These studies greatly improve our understanding of the evolution of human diseases. Nanopore sequencing technology also opens up new realms for epigenetic research, such as the telomere-to-telomere assembly and DNA methylation status of the human X chromosome (Miga et al., 2020) and the potential to simultaneously detect CpG methylation and chromatin accessibility (Lee et al., 2020). In addition, nanopore sequencing also enables more comprehensive and accurate analysis for the fragment size distribution of cell-free DNA and circulating tumor DNA in the plasma (Liu et al., 2020; Thirunavukarasu et al., 2021; Underhill, 2021; Yu et al., 2021). In the near future, when these research results are combined with multi-platform techniques and the analysis and mining of multi-omics data, early detection and early diagnosis of cancer and the detection of minimal residual disease in clinical practice will be more achievable. However, nanopore sequencing comes with obvious shortcomings, such as non-random system errors in its data and its over-reliance on base calling models (Liu et al., 2021; Wang et al., 2021), putting great demands on downstream analysis methods and tools. As a result, improving the accuracy of the sequencing and analysis results has been the greatest challenge facing nanopore sequencing.

NanoSquiggleVar is a method for direct analysis of targeted variant detection based on nanopore sequencing signals. It finds variant signals directly from the original sequencing signal without converting it to base sequences. NanoSquiggleVar avoids the need to build large and complex models and the loss of information due to the random nature of neural networks in the process of base calling and realizes targeted variant detection without the need of downstream bioinformatic analysis tools while still providing more accurate analysis results. For example, targeted point mutation detection and analysis on simulated NGS and nanopore data showed that both methods effectively located the expected mutation targets, with a consistency of 100% (Supplementary Sheet Table S3). However, if the detection was considered successful as long as one mutation was detected, then both G5230T in Sample-2 and A5648C in Sample-4 of the simulated NGS data were considered successful detections (Supplementary Figure S1). For the number of mutant reads detected, the deviation between the simulated nanopore results and the expected read number fluctuates more than that of NGS (Supplementary Figure S2). The p-values for the two were 0.66 and 4.23e−09, respectively, by two-tailed paired t test. These fluctuations could interfere with the test results in the actual sequencing, resulting in false positive and/or false negative results. NanoSquiggleVar does not produce these problems. And as we known, nanopore sequencing technology greatly expands the capacity of long-range, single-molecule DNA-modification detection, such as N6-methyladenine (6mA), N4-methylcytosine (4mC), and 5-methylcytosine (5mC) and its oxidative derivatives, i.e., 5-hydroxymethylcytosine (5hmC), 5-formylcytosine (5fC), and 5-carboxylcytosine (5caC). Nanopore sequencing detect DNA modifications via differences in the electric current intensity produced from a nanopore read of an unmodified base and that of a modified base. Therefore, NanoSquiggleVar is also theoretically suitable for the detection and analysis of targeted base modifications. Of course, we are trying our best to test and analyze this data with NanoSquiggleVar, and the results will be published in the future.

However, many shortcomings also exist in this novel method. The current analysis speed is very slow and requires considerable CPU resources and multithreaded parallel analysis. Targeted WSS and VSS signal sets need to be established. The simulated signal varies with the simulation software. For example, the signal values simulated by Scrappie and Deepsimulator showed substantial differences (data not shown), making it necessary to vary the DTW distance filtering threshold of NanoSquiggleVar according to the actual situation. NanoSquiggleVar can only determine the existence of a mutation and not its frequency. It can only be used when the number of mutation targets to be detected is small, and it cannot be applied on large genomes or to detect a large number of mutations. Much optimization work remains to be done on the algorithm to improve the operation efficiency. The method also needs to be adapted for use with nanopore sequencing instruments. These are the areas we need to address in the future, and we will make efforts to realize these improvements.

## CONCLUSIONS

NanoSquiggleVar is a method for the direct analysis of nanopore signals for targeted variant detection. It performs well on the detection and analysis of point mutations, indels, and homopolymer sequences. We hope NanoSquiggleVar can supplement the methodology of targeted variant detection by nanopore sequencing and promote the applications of nanopore real-time sequencing technology.

## Supporting information

Supplementary Sheet Table S1-S3

Supplementary Matrix

## DATA AVAILABILITY STATEMENT

The raw DTW distance value matrix of this study was all in the Supplementary Matrix-1 file and Supplementary Matrix-2 file.

## AUTHOR CONTRIBUTIONS

Jidong Lang designed the study, collected, simulated, analyzed and interpreted the data, and wrote the manuscript.

## References

Chen, Y., Zhou, X., and Yang, M. (2020). Nanopore Sequencing and Detection of Tumor Mutations. In Detection Methods in Precision Medicine, M. Yang, and M. Thompson, eds. (The Royal Society of Chemistry), p. 0.

Edge, P., and Bansal, V. (2019). Longshot enables accurate variant calling in diploid genomes from single-molecule long read sequencing. Nat Commun 10, 4660.

Goenka, S.D., Gorzynski, J.E., Shafin, K., Fisk, D.G., Pesout, T., Jensen, T.D., Monlong, J., Chang, P.C., Baid, G., Bernstein, J.A., et al. (2022). Accelerated identification of disease-causing variants with ultra-rapid nanopore genome sequencing. Nat Biotechnol 40, 1035–1041.

Katsman, E., Orlanski, S., Martignano, F., Fox-Fisher, I., Shemer, R., Dor, Y., Zick, A., Eden, A., Petrini, I., Conticello, S.G., et al. (2022). Detecting cell-of-origin and cancer-specific methylation features of cell-free DNA from Nanopore sequencing. Genome Biol 23, 158.

Kolmogorov, M., Yuan, J., Lin, Y., and Pevzner, P.A. (2019). Assembly of long, error-prone reads using repeat graphs. Nat Biotechnol 37, 540–546.

Koren, S., Walenz, B.P., Berlin, K., Miller, J.R., Bergman, N.H., and Phillippy, A.M. (2017). Canu: scalable and accurate long-read assembly via adaptive k-mer weighting and repeat separation. Genome Res 27, 722–736.

Lang, J. (2022). NanoCoV19: An analytical pipeline for rapid detection of severe acute respiratory syndrome coronavirus 2. Front Genet 13, 1008792.

Lang, J., Sun, J., Yang, Z., He, L., He, Y., Chen, Y., Huang, L., Li, P., Li, J., and Qin, L. (2022). Nano2NGS-Muta: a framework for converting nanopore sequencing data to NGS-liked sequencing data for hotspot mutation detection. NAR Genom Bioinform 4, lqac033.

Lang, J., Xu, Z., Wang, Y., Sun, J., and Yang, Z. (2023). NanoSTR: A method for detection of target short tandem repeats based on nanopore sequencing data. Front Mol Biosci 10, 1093519.

Lee, I., Razaghi, R., Gilpatrick, T., Molnar, M., Gershman, A., Sadowski, N., Sedlazeck, F.J., Hansen, K.D., Simpson, J.T., and Timp, W. (2020). Simultaneous profiling of chromatin accessibility and methylation on human cell lines with nanopore sequencing. Nat Methods 17, 1191–1199.

Li, H. (2018). Minimap2: pairwise alignment for nucleotide sequences. Bioinformatics 34, 3094–3100.

Li, H., and Durbin, R. (2009). Fast and accurate short read alignment with Burrows-Wheeler transform. Bioinformatics 25, 1754–1760.

Li, H., Handsaker, B., Wysoker, A., Fennell, T., Ruan, J., Homer, N., Marth, G., Abecasis, G., Durbin, R., and Genome Project Data Processing, S. (2009). The Sequence Alignment/Map format and SAMtools. Bioinformatics 25, 2078–2079.

Li, Y., Han, R., Bi, C., Li, M., Wang, S., and Gao, X. (2018). DeepSimulator: a deep simulator for Nanopore sequencing. Bioinformatics 34, 2899–2908.

Li, Y., Wang, S., Bi, C., Qiu, Z., Li, M., and Gao, X. (2020). DeepSimulator1.5: a more powerful, quicker and lighter simulator for Nanopore sequencing. Bioinformatics 36, 2578–2580.

Liu, X., Lang, J., Li, S., Wang, Y., Peng, L., Wang, W., Han, Y., Qi, C., Song, L., Yang, S., et al. (2020). Fragment Enrichment of Circulating Tumor DNA With Low-Frequency Mutations. Front Genet 11, 147.

Liu, Y., Rosikiewicz, W., Pan, Z., Jillette, N., Wang, P., Taghbalout, A., Foox, J., Mason, C., Carroll, M., Cheng, A., et al. (2021). DNA methylation-calling tools for Oxford Nanopore sequencing: a survey and human epigenome-wide evaluation. Genome Biol 22, 295.

Magi, A., Giusti, B., and Tattini, L. (2017). Characterization of MinION nanopore data for resequencing analyses. Brief Bioinform 18, 940–953.

Magi, A., Semeraro, R., Mingrino, A., Giusti, B., and D’Aurizio, R. (2018). Nanopore sequencing data analysis: state of the art, applications and challenges. Brief Bioinform 19, 1256–1272.

Martignano, F., Munagala, U., Crucitta, S., Mingrino, A., Semeraro, R., Del Re, M., Petrini, I., Magi, A., and Conticello, S.G. (2021). Nanopore sequencing from liquid biopsy: analysis of copy number variations from cell-free DNA of lung cancer patients. Mol Cancer 20, 32.

Miga, K.H., Koren, S., Rhie, A., Vollger, M.R., Gershman, A., Bzikadze, A., Brooks, S., Howe, E., Porubsky, D., Logsdon, G.A., et al. (2020). Telomere-to-telomere assembly of a complete human X chromosome. Nature 585, 79–84.

Mitsuhashi, S., Frith, M.C., Mizuguchi, T., Miyatake, S., Toyota, T., Adachi, H., Oma, Y., Kino, Y., Mitsuhashi, H., and Matsumoto, N. (2019). Tandem-genotypes: robust detection of tandem repeat expansions from long DNA reads. Genome Biol 20, 58.

Rang, F.J., Kloosterman, W.P., and de Ridder, J. (2018). From squiggle to basepair: computational approaches for improving nanopore sequencing read accuracy. Genome Biol 19, 90.

Shafin, K., Pesout, T., Chang, P.C., Nattestad, M., Kolesnikov, A., Goel, S., Baid, G., Kolmogorov, M., Eizenga, J.M., Miga, K.H., et al. (2021). Haplotype-aware variant calling with PEPPER-Margin-DeepVariant enables high accuracy in nanopore long-reads. Nat Methods 18, 1322–1332.

Shen, W., Le, S., Li, Y., and Hu, F. (2016). SeqKit: A Cross-Platform and Ultrafast Toolkit for FASTA/Q File Manipulation. PLoS One 11, e0163962.

Tarasov, A., Vilella, A.J., Cuppen, E., Nijman, I.J., and Prins, P. (2015). Sambamba: fast processing of NGS alignment formats. Bioinformatics 31, 2032–2034.

Teng, H., Cao, M.D., Hall, M.B., Duarte, T., Wang, S., and Coin, L.J.M. (2018). Chiron: translating nanopore raw signal directly into nucleotide sequence using deep learning. Gigascience

Thirunavukarasu, D., Cheng, L.Y., Song, P., Chen, S.X., Borad, M.J., Kwong, L., James, P., Turner, D.J., and Zhang, D.Y. (2021). Oncogene Concatenated Enriched Amplicon Nanopore Sequencing for rapid, accurate, and affordable somatic mutation detection. Genome Biol 22, 227.

Underhill, H.R. (2021). Leveraging the Fragment Length of Circulating Tumour DNA to Improve Molecular Profiling of Solid Tumour Malignancies with Next-Generation Sequencing: A Pathway to Advanced Non-invasive Diagnostics in Precision Oncology? Mol Diagn Ther 25, 389–408.

Wan, Y.K., Hendra, C., Pratanwanich, P.N., and Goke, J. (2022). Beyond sequencing: machine learning algorithms extract biology hidden in Nanopore signal data. Trends Genet 38, 246–257.

Wang, Y., Zhao, Y., Bollas, A., Wang, Y., and Au, K.F. (2021). Nanopore sequencing technology, bioinformatics and applications. Nat Biotechnol 39, 1348–1365.

Wick, R.R., Judd, L.M., and Holt, K.E. (2019). Performance of neural network basecalling tools for Oxford Nanopore sequencing. Genome Biol 20, 129.

Yu, S.C.Y., Jiang, P., Peng, W., Cheng, S.H., Cheung, Y.T.T., Tse, O.Y.O., Shang, H., Poon, L.C., Leung, T.Y., Chan, K.C.A., et al. (2021). Single-molecule sequencing reveals a large population of long cell-free DNA molecules in maternal plasma. Proc Natl Acad Sci U S A 118.

Garrison, E. and Marth., G. (2012). Haplotype-based variant detection from short-read sequencing. 2012:1207.3907v2. (preprint: not peer reviewed)

