## Supplementary figures and images for "NanoSquiggleVar: A method for direct analysis of targeted variants based on nanopore sequencing signals"

### Supplementary Figure S1

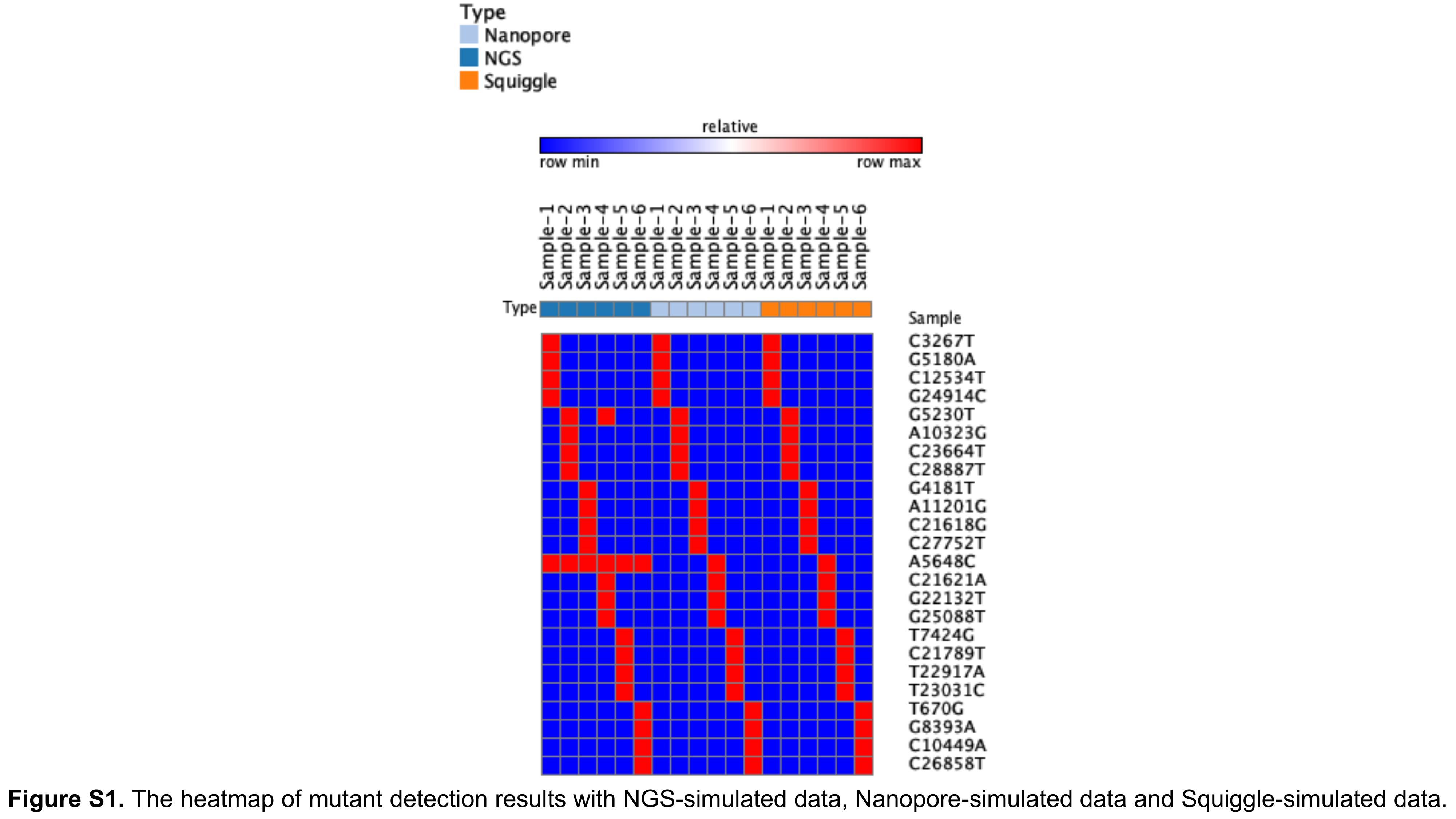

### Supplementary Figure S2

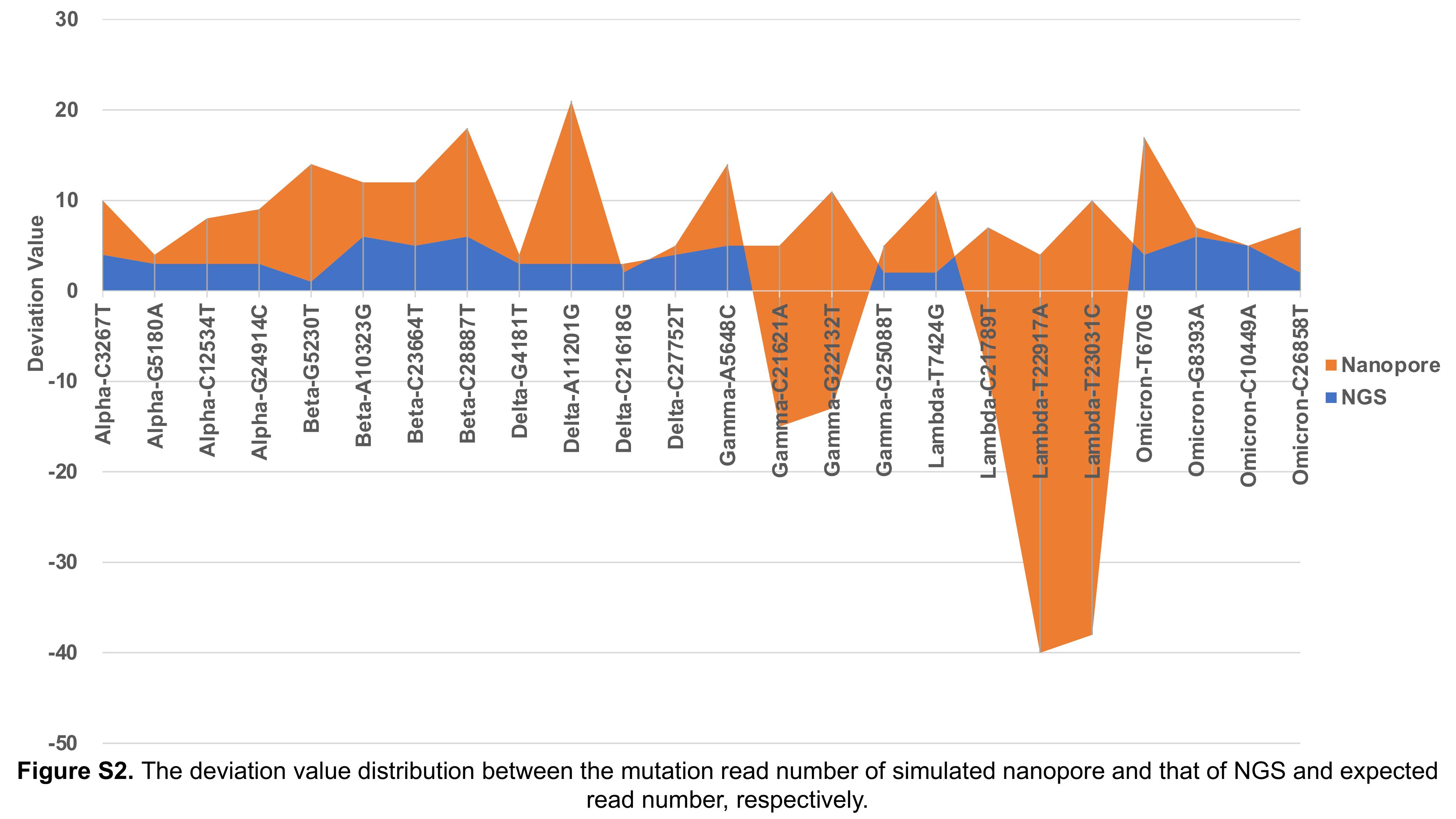
